# Human-genome gut-microbiome interaction in Parkinson’s disease

**DOI:** 10.1101/2021.01.21.427679

**Authors:** Zachary D. Wallen, William J. Stone, Stewart A. Factor, Eric Molho, Cyrus P. Zabetian, David G. Standaert, Haydeh Payami

## Abstract

The causes of complex diseases remain an enigma despite decades of epidemiologic research on environmental risks and genome-wide studies that have uncovered tens or hundreds of susceptibility loci for each disease. We hypothesize that the microbiome is the missing link. Genetic studies have shown that overexpression of alpha-synuclein, a key pathological protein in Parkinson’s disease (PD), can cause familial PD and variants at alpha-synuclein locus confer risk of idiopathic PD. Recently, dysbiosis of gut microbiome in PD was identified: altered abundances of three microbial clusters were found, one of which was composed of opportunistic pathogens. Using two large datasets, we show that the overabundance of opportunistic pathogens in PD gut is influenced by the host genotype at the alpha-synuclein locus, and that the variants responsible modulate alpha-synuclein expression. This is the first demonstration of interaction between genetic factors in the human genome and the dysbiosis of gut microbiome in PD.

## Introduction

Parkinson’s disease (PD) affects over 6 million people world-wide, having doubled in one decade, and continues to rapidly increase in prevalence with the aging of the world population^1^. PD is a progressive degenerative disease which affects the brain, the peripheral nervous system, and the gastrointestinal tract, causing progressive, debilitating movement disorders, gastrointestinal and autonomic dysfunction, sleep disorders, and cognitive impairment. Currently there is no prevention, cure or treatment known to slow the progression of the disease.

Like other common late-onset disorders, PD has Mendelian forms caused by rare mutations, but the vast majority of cases remain idiopathic. Both genetic and environmental risk factors have been identified^2–4^, but none have large enough effect sizes individually or in combination to fully encapsulate disease risk^5–8^. The triggers that initiate onset of PD pathology are unknown.

There is a connection between PD and the gastrointestinal tract^9,10^ and the gut microbiome^11^. The gut microbiome is a relatively new and increasingly active area of research in human disease^12–14^. Studies on PD have consistently found altered gut microbiome, with depletion of short-chain fatty acid (SCFA) producing bacteria, and enrichment of *Lactobacillus* and *Bifidobacterium*^11,15,16^. Most studies to date have been modest in size, and therefore have examined mostly common microorganisms. No study so far has had the power to explore interactions between gut microorganisms and genetic risk factors for PD. Demonstrating interaction is statistically challenging because data are parsed into smaller groups, which drastically reduces effective sample size and power. We recently reported a microbiome-wide association study in PD, using two large datasets and internal replication, which enabled investigation of less common taxa not reported before^11^. In these datasets, reduced SCFA-producing bacteria and elevated *Lactobacillus* and *Bifidobacterium* were robustly confirmed. In addition, a significant increase was detected in the relative abundance of a poly-microbial cluster of opportunistic pathogens, including *Corynebacterium_1 (C. amycolatum, C. lactis), Porphyromonas (P. asaccharolytica, P. bennonis, P. somerae, P. uenonis*), and *Prevotella (P. bivia, P. buccalis, P. disiens, P. timonensis*). These are commensal bacteria with normally low abundance in the gut, but they can cause infections in opportunistic situations such as a compromised immune system^11^.

Overabundance of opportunistic pathogens in PD gut was of interest because it harks back to the hypothesis advanced by Professor Heiko Braak which proposes that in non-familial forms of PD, the disease is triggered by an unknown pathogen in the gut and spreads to the brain^17,18^. Braak’s hypothesis was based on pathological studies of postmortem human brain, stained using antibodies to alpha-synuclein. Misfolded alpha-synuclein, the pathologic hallmark of PD, has been seen to form in enteric neurons early in disease^19–21^, and has been shown to propagate in a prion-like manner from the gut to the brain in animal models^22^. The gene that encodes alpha-synuclein is *SNCA. SNCA* gene multiplication results in drastic over expression of alpha-synuclein and causes Mendelian autosomal dominant PD. Variants in the *SNCA* region are associated with risk of idiopathic PD^23^, and are expression quantitative trait loci (eQTL) associated with expression levels of *SNCA*^24–26^. Increased alpha-synuclein expression has been noted with infections unrelated to PD^27,28^. We hypothesized that if opportunistic pathogens are involved in disease pathogenesis, there might be an interaction between genetic variants in *SNCA* region and dysbiosis of the gut in PD.

## Results

The two case-control cohorts used here are those previously employed by Wallen et al. to characterize the PD gut microbiome^11^. Here, we generated and added genotype data to investigate interactions. The sample size for the present analysis was 199 PD and 117 controls in dataset 1, and 312 PD and 174 controls in dataset 2. All samples had complete genotypes, 16S microbiome data, and metadata (Supplementary Table 1).

We defined the boundaries of the *SNCA* region such that it would encompass known *cis*-eQTLs for *SNCA*. Using GTEx eQTL database, we defined the boundaries as ch4:88.9Mb, downstream of 3’ *SNCA*, and ch4:90.6 Mb, upstream of 5’ *SNCA*. In our genome-wide genotype data (see Methods), we had 2,627 single nucleotide polymorphisms (SNPs) that mapped to this region, had minor allele frequency (MAF) >0.1, imputation quality score *r^2^*>0.8, and were in common between the two datasets being studied here.

The taxa examined were grouped and analyzed at genus/subgenus/clade level as *Corynebacterium_1 (C. amycolatum, C. Lactis), Porphyromonas (P. asaccharolytica, P. bennonis, P. somerae, P. uenonis*), and *Prevotella (P. bivia, P. buccalis, P. disiens, P. timonensis*). For simplicity, we will refer to the three microbial groups as taxa. As we have previously shown, the abundance of these taxa are elevated in PD vs. control. These findings were replicated in the two datasets (Table 1), verified by two statistical methods, robust to covariate adjustment (over 40 variables investigated), and yielded no evidence of being the result of PD medications or disease duration^11^.

**Table 1.**
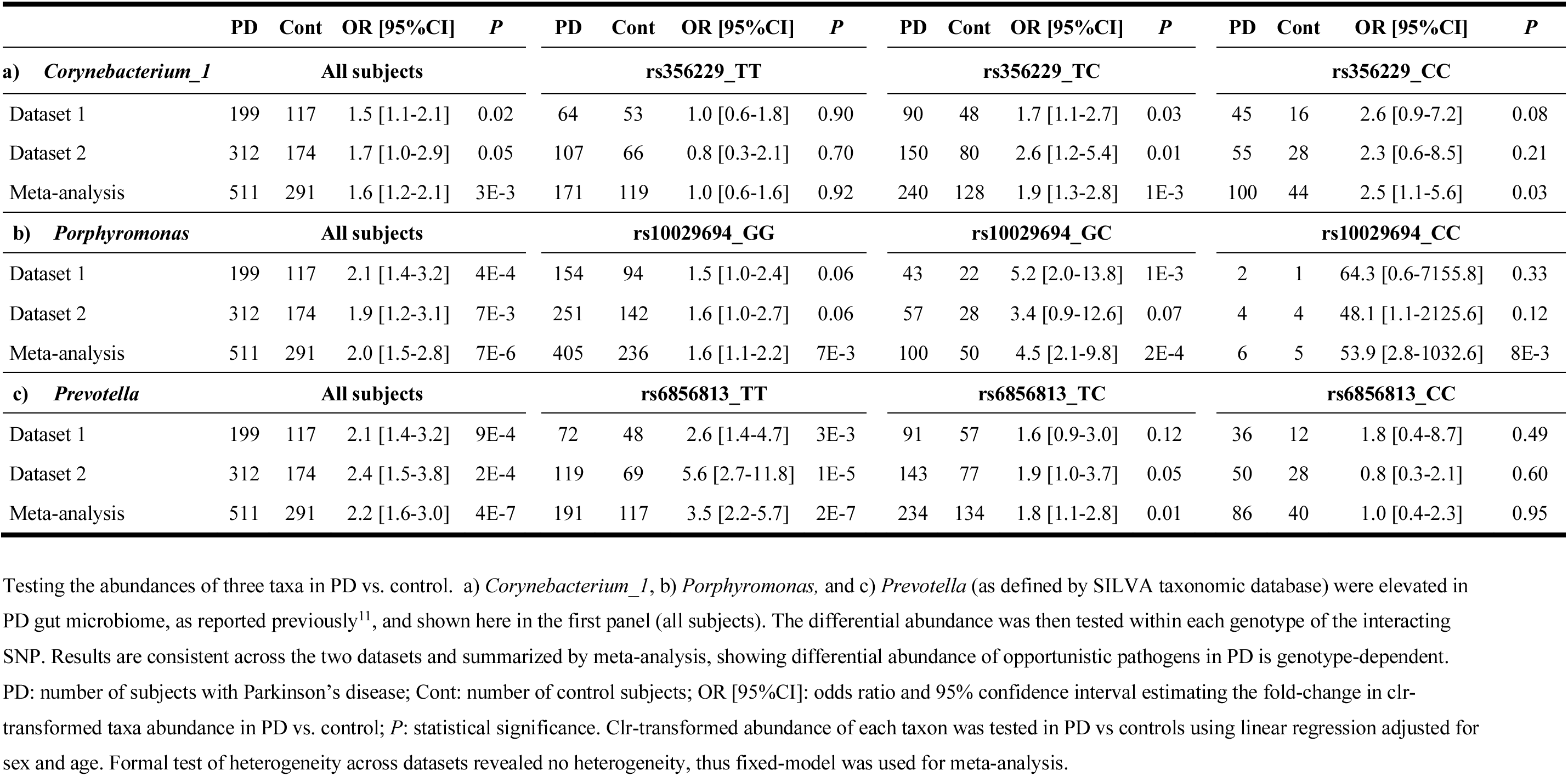
Increased abundance of opportunistic pathogens in PD gut microbiome is dependent on the host genotype.

The analysis of interaction was structured as follows. (1) We screened for statistical interaction between 2,627 SNP genotypes in the *SNCA* region, case-control status, and centered log-ratio (clr) transformed abundance of each taxon, and selected the SNP with the highest statistical significance as the candidate interacting SNP (Fig. 1a-c). (2) We then tested association of each taxon with case-control status after stratifying the subjects by the interacting SNP genotype. The effect of SNP on PD-taxa association was tested statistically (Table 1) and illustrated graphically (Fig. 2, Supplementary Fig. 1). (3) We tested association of interacting SNPs with PD (Table 2). This test was conducted because interaction can exist with or without a main effect of SNP on disease risk. SNPs with a main effect are detected in GWAS, modifiers without a main effect are missed in GWAS^5,7^. (4) We conducted *in silico* functional analysis of the interacting SNPs (Table 2, Fig. 1d,e). All analyses were performed in two datasets, followed-by meta-analysis.

**Fig. 1:**
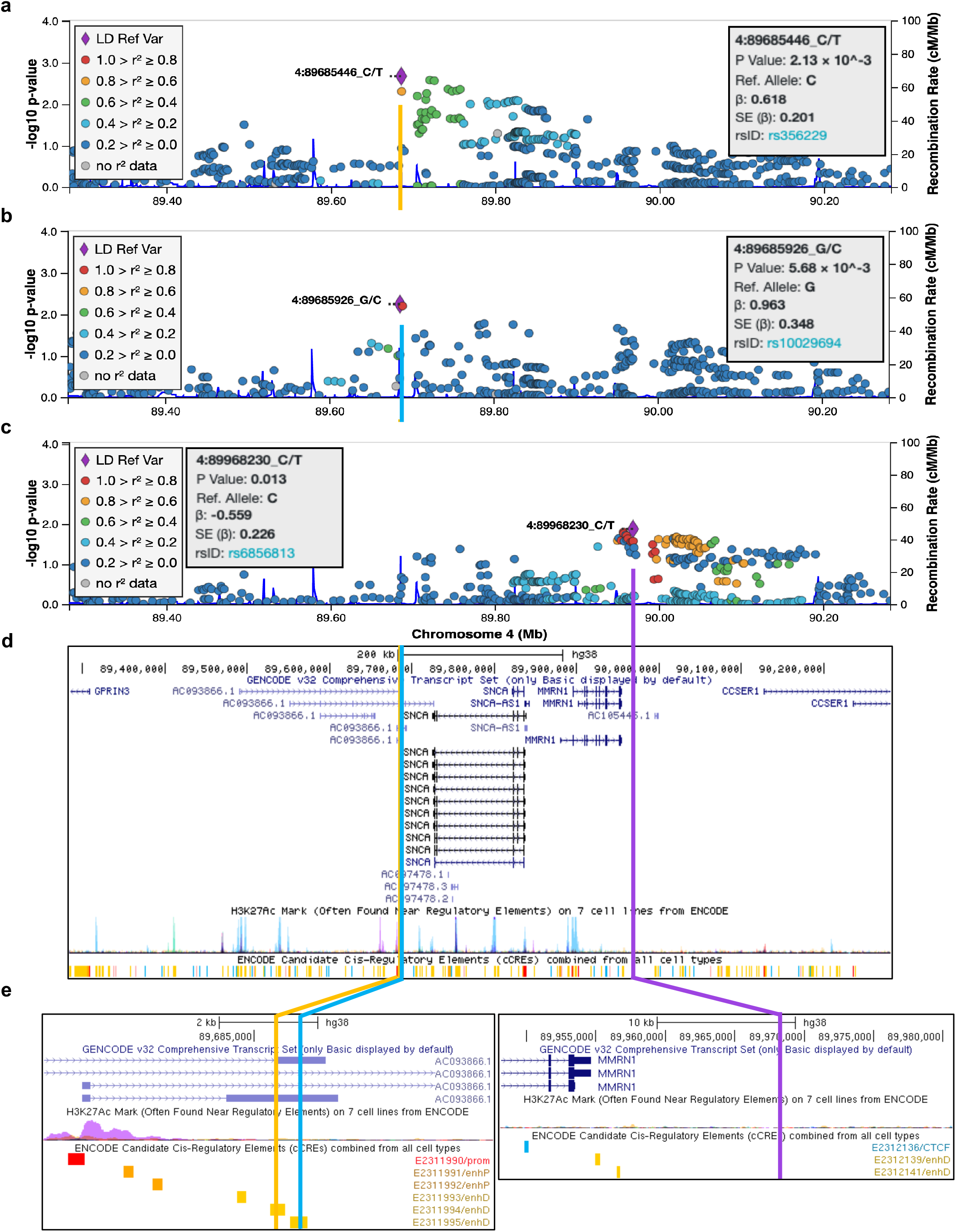
Genetic map of candidate interacting SNPs. SNPs in *SNCA* region (chromosome 4: 88.9 Mb – 90.6 Mb) were tested for interaction on the associaion of three taxa with PD. Results are shown in LocusZoom, where each SNP is plotted according to its base pair position and meta-analysis -log10(*P* value) for interaction for the three taxa: (a) *Corynebacterium_1*, (b) *Porphyromonas*, and (c) *Prevotella*. The SNP with the highest significance is shown as a purple diamond, and was chosen as candidate interacting SNP for stratified analysis (Table 1). (d) UCSC Genome Browser shows the interacting SNPs for *Corynebacterium_1* and *Porphyromonas* map to 3’ *SNCA* in a lncRNA that overlaps with and are antisense to *SNCA*. The interacting SNP for *Prevotella* is distal at 5’ of *SNCA* and *MMRN1*. (e) The interacting SNPs for *Corynebacterium_1* and *Porphyromonas*, while only 450 base pair apart, are not in LD (R^2^=0) and map to adjacent regulatory sequences shown in yellow bars. The interacting SNP for *Prevotella* does not map to any known functional sequence. All three SNPs are eQTLs for *SNCA* and lncRNA genes *SNCA-AS1, RP11-115D19.1 (AC093866.1*), and *RP11-115D19.2 (AC097478.2*) which are associated with expression of *SNCA* (Table 2). LD: linkage disequilibrium; Mb: Megabase; P value: *P* value from meta-analysis; β: beta coefficient of interaction from meta-analysis; SE: standard error; rsID: reference SNP ID for the marked SNP.

**Fig. 2:**
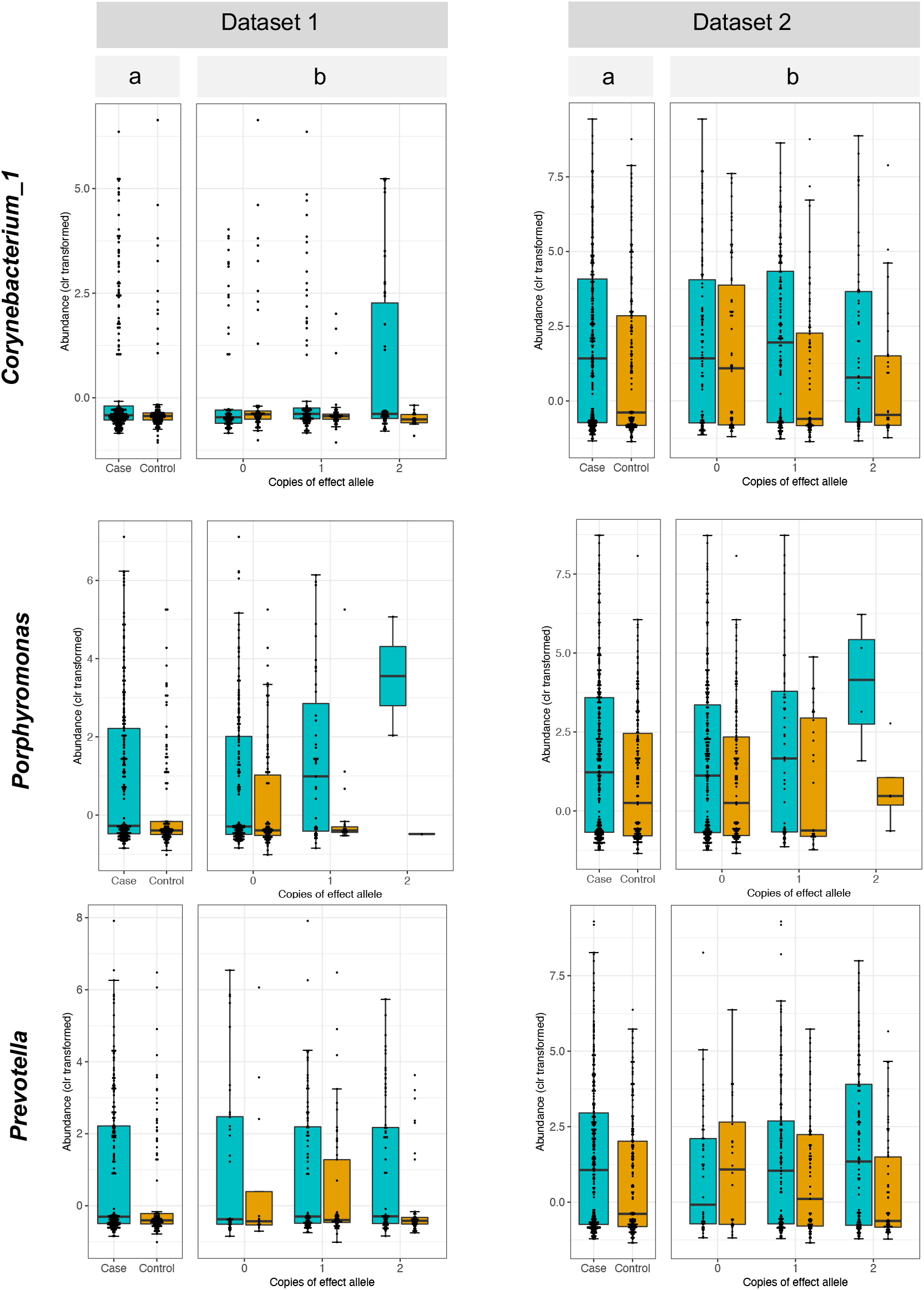
Differential abundance of opportunistic pathogens. Clr-transformed abundances of each taxon is plotted for PD cases (blue) and controls (orange) for all subjects irrespective of genotype (panel a) and stratified for the three genotypes of the interacting SNP (panel b). The two datasets show the same pattern of interaction where the difference between PD and controls in the abundances of each taxon becomes larger with increasing number of the effect allele. Dataset 2 has higher resolution than dataset 1 (particularly for *Corynebacterium_1* which is rare) because it had 10x greater sequencing depth.

**Table 2.**
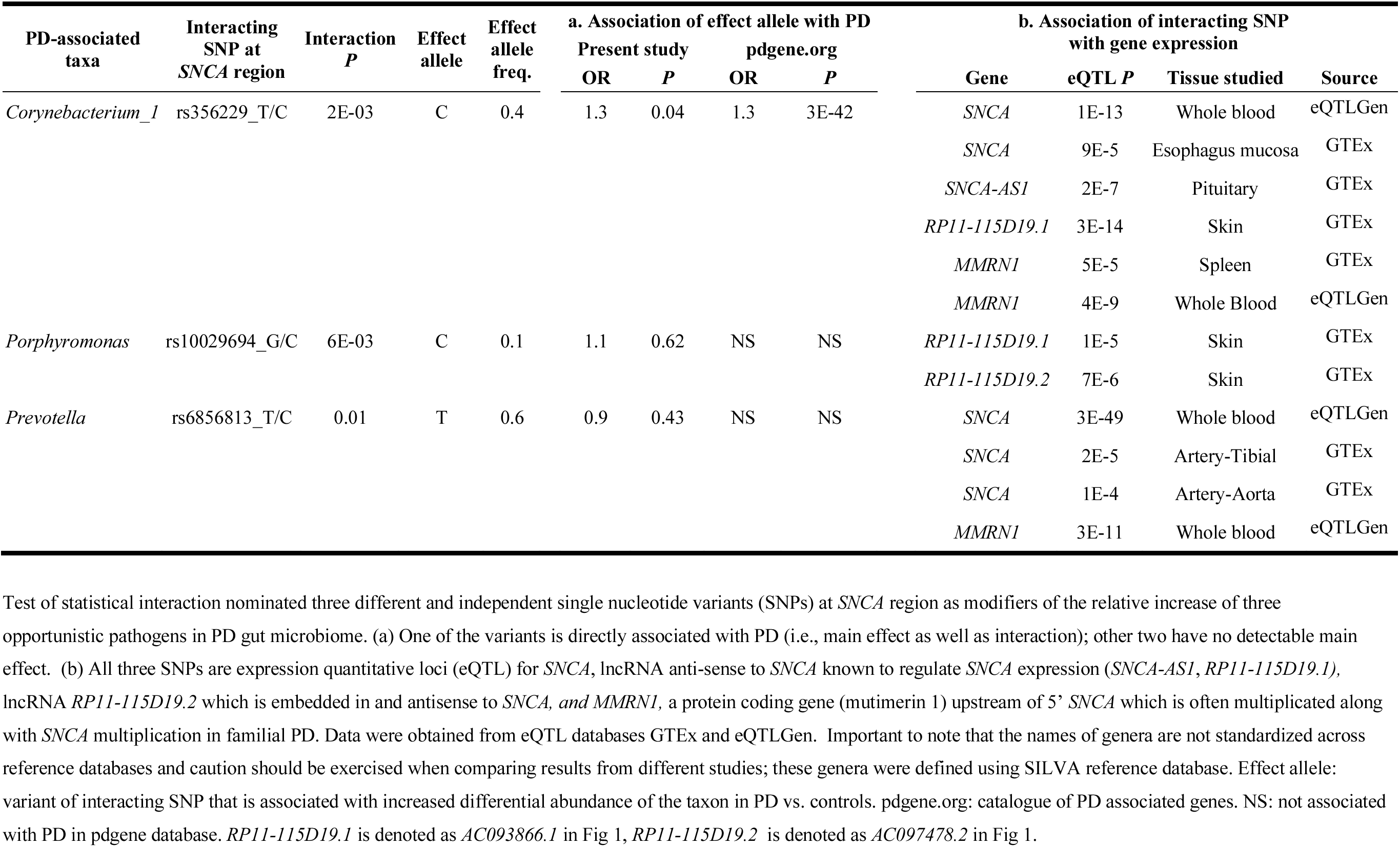
Characteristics of the interacting variants at *SNCA* locus

### Corynebacterium_1

1. *Screening for interaction* (Fig. 1a). The candidate interacting SNP for *Corynebacterium_1* was rs356229 (interaction *P*=2E-3). rs356229 is located 3’ of *SNCA* (Fig. 1). The two alleles are rs356229_T (allele frequency=0.6) and rs356229_C (frequency=0.4). rs356229 was imputed with imputation quality score of 0.96 in dataset 1 and 0.99 in dataset 2.
2. *PD-taxa association varied by genotype* (Table 1a, Fig. 2). If we do not consider genotype, *Corynebacterium_1* abundance is significantly elevated in PD (OR=1.6, *P*=3E-3). However, when data are stratified by genotype, there is no association between *Corynebacterium_1* and PD among individuals with rs356229_TT genotype, who comprised 36% of the study (OR=1.0, *P*=0.92). The association of *Corynebactreium_1* with PD was dependent on the presence of the rs356229_C allele. The abundance of *Corynebactreium_1* was nearly 2-fold higher in PD than controls in heterozygous rs356229_CT (OR=1.9, *P*=1E-3), and 2.5-fold higher in the homozygous rs356229_CC individuals (OR=2.5, *P*=0.03).
3. *Association of SNP with PD* (Table 2). rs356229 has been previously identified in PD GWAS meta-analysis, with the rs356229_C allele associated with increased PD risk (OR=1.3, *P*=3E-42 with N=108,990 samples, pdgene.org^23^). We also detected an association between rs356229_C and PD in the present dataset (OR=1.3, *P*=0.04 with N=802 samples). That we estimated an effect size identical to GWAS, despite the enormous disparity in the sample size and power, speaks to the robustness of the data. Interestingly, the association of rs356229_C with risk of PD varied by the increasing abundance of *Corynebacterium_1* from no association in the 1^st^ or 2^nd^ quartile (OR=0.9, *P*=0.5; OR=1.1, *P*=0.8) to an emerging and then strong association in the 3^rd^ and 4^th^ quartiles (OR=1.4, *P*=0.1 and OR=2.2, *P*=5E-3). The interactive effect of rs356229 on the association of *Corynebacterium_1* and PD does not stem from its direct association with PD. This can be seen in Table 1a, where the test is between *Corynebacterium_1* and PD; rs356229 is not in the test, it was only used to divide the samples by genotype, which showed varying association between the taxon and PD as a function of genotypes. Also of note is that rs356182, which is the highest SNP in the *SNCA* peak in PD GWAS (OR=1.34, *P*=2E-82), yielded no evidence for interaction with the taxa tested here, underscoring the notion that the variants that show the strongest association with disease are not necessarily the best candidates for interaction.
4. *Functional analysis in silico* (Fig. 1d,e, Table 2). rs356229 maps to a distal regulatory element at 3’ of *SNCA*. rs356229 is an eQTL for *SNCA*. Data were obtained by eQTL GWAS conducted in whole blood (eQTLGen.org) and in esophagus mucosa (GTExportal.org). rs356229_C allele is associated with increased expression of *SNCA* in blood (eQTL *P*=1E-13) and in esophagus mucosa (eQTL *P*=9E-5). According to GTEx, rs356229 is also an eQTL for *SNCA-AS1* (eQTL *P*=2E-7) and *RP11-115D19.1* (eQTL *P*=3E-14). *SNCA-AS1* and *RP11-115D19.1* overlap with *SNCA* and encode long non-coding RNA (lncRNA) that are antisense to *SNCA* (Fig. 1d) and have been implicated in regulation of *SNCA* expression^29–31^.

### Porphyromonas

1. *Screening for interaction* (Fig. 1b). The candidate interacting SNP for *Porphyromonas* was rs10029694 (interaction *P*=6E-3). rs10029694 maps to 3’ of *SNCA* (Fig. 1). The two alleles are rs10029694_G (frequency=0.9) and rs10029694_C (frequency=0.1). rs10029694 was imputed with imputation quality score 0.99 in dataset 1 and 0.92 in dataset 2. The interacting SNPs for *Porphyromonas* (rs10029694) and *Corynebacterium_1* (rs356229) map very close to each other, only 480 base pairs apart, but they are not in linkage disequilibrium (LD): D’<0.01, R^2^=0.
2. *PD-taxa association varies by genotype* (Table 1b, Fig. 2). *Porphyromonas* was elevated in PD irrespective of rs10029694_G/C genotype (OR=2.0, *P*=7E-6), and in every genotype, but the statistical interaction implied difference across genotypes. Shown in stratified analysis (Table 1b), the rs10029694_GG genotype had a nearly two-fold higher abundance of *Porphyromonas* in PD vs. controls (OR=1.6, P=7E-3), rs10029694_GC had nearly five-fold difference (OR=4.5, P=2E-4) and rs10029694_CC had approximately 54-times higher abundance of *Porphyromonas* in PD than in controls (OR=53.9, P=8E-3). Note however that there were only 11 individuals with rs10029694_CC genotype. Although the statistical methods were carefully chosen to be robust to small sample size, and the *P* value is quite significant despite the sample size, the fact remains that the OR=54 was generated on only 11 people. If we collapse the rare rs10029694_CC genotype with rs10029694_CG, we have 161 individuals (20% of subjects) with at least one copy of rs10029694_C allele, and we get a more conservative estimate of OR=5.1 (*P*=2E-5) for association of *Porphyromonas* with PD in people with one or two copies of rs10029694_C.
3. *Association of SNP with PD* (Table 2). rs10029694 has not been nominated by GWAS as a risk variant for PD, nor does it show evidence for association with PD in our datasets (OR=1.1, *P*=0.6). This variant appears to impart an effect on the association of *Porphyromonas* with PD without having a main effect on PD. There are published examples of modifiers (*GRIN2A, SV2C*) that had no detectable main effect in GWAS but were found through interaction (with caffeine use and smoking) and were subsequently shown experimentally to play key roles in PD pathogenesis^5,7^. As would be expected from the interaction, the frequency of the effect allele rs10029694_C in PD vs. control rose with increasing abundance of *Porphyromonas*, yielding OR=0.6 (*P*=0.2) for 1^st^ quartile and increasing up to OR=2.2 (*P*=0.08) for the 4^th^ quartile.
4. *Functional analysis in silico* (Fig. 1d,e and Table 2). rs10029694 maps to a distal regulatory element at 3’ of *SNCA*, adjacent to another regulatory element where rs356229, the interacting SNP for *Corynebacterium_1* resides. rs10029694 is an eQTL for two lncRNA that are antisense to *SNCA: RP11-115D19.1* (eQTL *P*=1E-5) and *RP11-115D19.2* (eQTL *P*=7E-6). *RP11-115D19.1* overlaps with 3’ of *SNCA; RP11-115D19.2* is within *SNCA*. We did not find direct evidence for rs10029694 being an eQTL for *SNCA*. However, *RP11-115D19.1* and *RP11-115D19.2* are anti-sense to *SNCA* which based on current knowledge on function of antisense lncRNA would be presumed to be regulatory for *SNCA*^30,31^, and *RP11-115D19.1* has been directly shown to regulate *SNCA* expression^29^.

### Prevotella

1. *Screening for interaction* (Fig. 1c). The candidate interacting SNP for *Prevotella* was rs6856813 (interaction *P*=0.01). rs6856813 is ~100kb upstream at 5’ of *SNCA*. The two alleles are rs6856813_T (frequency=0.6) and rs6856813_C (frequency=0.4). rs6856813 was imputed with imputation quality score 0.98 in dataset 1 and 0.84 in dataset 2. rs6856813 is 300Kb away from and not in LD with the interacting SNPs of *Corynebacterium_1* (rs356229, D’= 0.2, R^2^=0.04) or *Porphyromonas* (rs10029694, D’=0.36, R^2^=0.01).
2. *PD-taxa association varies by genotype* (Table 1c, Fig. 2). *Prevotella* was elevated two-fold in PD vs. controls (OR=2.2, *P*=4E-7). Genotype-specific results suggest rs6856813_TT had the greatest differential abundance in PD vs. control (OR=3.5, *P*=2E-7), followed by rs6856813_TC (OR=1.8, *P*=0.01), and no difference in rs6856813_CC genotype (OR=1.0, *P*=0.95).
3. *Association of SNP with PD* (Table 2). rs6856813_C/T had no main effect for association with PD in this study (OR=0.9, *P*=0.4) nor in PD GWAS^23^. There is a statistically non-significant trend of increasing frequency of rs6856813_T allele with increasing abundance of *Prevotella* in PD, yielding OR=0.8 in 1^st^ quartile and increasing to OR=1.5 in 4^th^ quartile, consistent with the presence of interaction.
4. *Functional analysis in silico* (Fig. 1d,e, Table 2). Although rs6856813 is ~100kb upstream of S*NCA* and does not map to a known regulatory sequence, it is a strong eQTL for *SNCA:* the rs6856813_T allele, which is the effect allele for interaction with *Prevotella*, is associated with increased *SNCA* expression in blood (eQTL *P*=3E-49) and in arteries (eQTL *P*=2E-5).

## Discussion

Numerous studies have been performed on the association of genetic variants with PD and separately of gut microbiome and PD, but this is the first, to our knowledge, that has attempted to study the interaction between the two. Here we have used a candidate taxa, candidate gene strategy: we used prior knowledge of the association of PD with elevated abundances of certain opportunistic pathogens in the gut^11^ and searched for genetic modifiers of these associations in the *SNCA* gene region^23^. Through statistical interaction tests we identified specific variants in the *SNCA* region as candidate interacting variants and through genotype-stratified analyses we showed that the increases in relative abundance of opportunistic pathogens in PD gut is modulated by host genotype.

Statistical interaction tests provide a means to investigate if association of one factor with the trait is influenced by a second factor. Interaction studies require much larger sample sizes and power than association studies; the P values for interaction seldom achieve significance, and when they do, they are far less significant that the P values for a similarly sized one-factor association study. Here, we tested if association of three opportunistic pathogens with PD (organisms with higher relative abundance in PD cases than similarly aged controls) is dependent on genetic variations in or around S*NCA*. Sometimes the interacting variant discovered by this approach is also directly associated with trait, but not always. In the present study, the SNP that affects the association of *Corynebacterium_1* with PD is also directly associated with PD (it is one of the SNPs in the *SNCA* peak in PD GWAS); in contrast, the interacting SNPs for *Porphyromonas* and *Prevotella* have no main effect that can be detected as association with PD. Factors that do not have a main effect on the trait are missed in association tests (e.g., GWAS). Thus, interaction testing is complementary to association testing in that it can identify novel markers that are otherwise missed. Two prior examples of PD-relevant genes that were missed by GWAS are synaptic vesicle 2C (*SV2C*) gene which emerged in interaction with smoking^7^ and led to deciphering its role in dopamine release and its disruption in PD^32^, and the gene encoding Glutamate Ionotropic Receptor NMDA Type Subunit 2A (*GRIN2A*) which was detected via interaction with caffeine intake^5^. Neither *SV2C* nor *GRIN2A* has a main effect on PD and were both missed in GWAS. The present study is conceptually similar to the *SV2C* and *GRIN2A* studies, but on a smaller scope because of the limited sample size. Interaction studies that revealed *SV2C* and *GRIN2A* were conducted on a genome-wide level with a sample size of approximately 1500 PD and 1500 controls for whom both genotype and smoking/caffeine data were available. The largest PD datasets that have both genotype and microbiome data are the two datasets used here, one has 199 PD and 117 controls and the other 312 PD and 174 controls. It will be important to collect larger datasets which will allow the exploration of genotype-microbiome interactions at the genome-wide and microbiome-wide level.

Our rationale for choosing *SNCA* and opportunistic pathogen as our candidate gene and candidate taxa stemmed from the collective literature. *SNCA* is a key player in PD. Alpha-synuclein aggregates are a pathologic hallmark of PD. Mutations in *SNCA* cause autosomal dominant PD and variants that affect *SNCA* gene expression are the most significant genetic risk factors for idiopathic PD^23,33^. While the functions of alpha-synuclein is yet to be fully understood, it has been shown to play a key role in activating the immune system, acting as antigen presented by PD-associated major histocompatibility molecules and recognized by T cells which infiltrate the brain^34–36^. *SNCA* expression has also been shown to be critical for inducing immune response against infections unrelated to PD^27,28^. Alpha-synuclein aggregates, which have historically been considered as a marker of PD pathology in the brain, can actually form in the enteric neurons^19^ and in animal models have been shown to propagate from the gut to the brain^22^ possibly via the vagus nerve^37,38^. The trigger that induces alpha-synuclein pathology in the gut is unknown. Braak hypothesized the trigger is a pathogen^17,18^. Our choice of opportunistic pathogens as the candidate taxa for interaction testing was driven by our recent finding of an overabundance of opportunistic pathogens in PD gut and Braak’s hypothesis. Moreover, a study conducted in mice has corroborated that intestinal infection triggers dopaminergic cell loss and motor impairment in a *Pink1* knockout model of PD^39^. Whether the opportunistic pathogens found in human PD microbiome are the triggers of PD is being investigated. In the meantime, we thought that if these opportunistic pathogens are involved in PD pathogenesis, there is likely a connection to *SNCA* genotype worth testing.

Interestingly, three different *SNCA*-linked genetic variants emerged as modifiers for the association of the three opportunistic pathogens with PD. They are independent of each other with no LD among them. All three interacting variants are eQTLs for *SNCA* and lncRNAs that affect expression of *SNCA*. This suggests a link between *SNCA* expression and presence of opportunistic pathogens, and that regulation of this link may involve different regulatory elements depending on the pathogen. We do not know if this is because of tissue specificity of gene expression. It is not known which cells in the gut are responsible for expression and corruption of alpha-synuclein into pathologic species. If the opportunistic pathogens induce *SNCA* expression, they may do so by signaling different cell types, hence the involvement of different regulatory elements. *Prevotella* and *Porphyromonas* are commensal to gastrointestinal and urinary track, *Corynebacterium* is common in skin microbiome. All three can be found at low abundance in the gut. All three have been implicated in causing infections in nearly every type of tissue (reviewed by Wallen et al.^11^).

These data provide new leads that with follow-up will yield a better understanding of disease pathogenesis. These data alone cannot resolve cause and effect. We cannot tell if the *SNCA* genotype leads to altered colonization of the gut, which in turn leads to PD, or is it the other way around, *SNCA* genotype causes PD (unlikely in the absence of a main effect on PD), which leads to gut dysfunction and accumulation of pathogens. Or, maybe the pathogen induces alpha-synuclein expression which elicits immune response to infection as seen in other infections unrelated to PD, but in individuals with certain regulatory genotypes at *SNCA*, alpha-synuclein expression goes into overdrive and PD is a down-stream consequence. Further studies in humans conducted over time and in experimental models will be needed to tease out the underlying biology of these interactions.

This study serves as proof of principal that genetic susceptibility to disease and the dysbiosis in the gut microbiome are not operating independently. Rather, it suggests that alterations in gut microbiome should be integrated in the gene-environment interaction paradigm, which has long been suspected to be the cause of idiopathic disease but is yet to produce a causative combination. To advance these ideas further, the biggest challenge is to secure well-coordinated studies with large sample sizes. Unlike genetic studies which can be pooled thanks to the stability of DNA, pooling of microbiome studies should be avoided due to effects of collection and storage parameters on outcomes. Standardization of methods can alleviate some of the cross-study variations. It is also more difficult to collect stool samples than, for example, smoking data or saliva. People are averse to donating stool samples; 30% of our research participants who donated blood refused to donate stool. Researchers are cognizant of the need to join resources, create standardized protocols, and coordinate data collection across laboratories. Within a few years, we will be able to amass the sample sizes needed to address interaction of genes, environment, and microbiome on a comprehensive scale.

## Methods

### Subjects

The study was approved by the institutional review boards at all participating institutions, namely New York State Department of Health, University of Alabama at Birmingham, VA Puget Sound Health Care System, Emory University, and Albany Medical Center. All subjects provided written informed consent for their participation. This study included two datasets each composed of persons with PD (case) and neurologically healthy individuals (control). Subject enrollment and data collection for both datasets was conducted by the NeuroGenetics Research Consortium (NGRC) team using uniform protocols. The two datasets used here were the same datasets used by Wallen et al for characterizing the microbiome^11^; except here we have generated and added genetic data, and subjects without genotype were excluded (Supplementary Table 1). Methods of subject selection and data collection have been described in detail before^11^. Briefly, PD was diagnosed by NGRC-affiliated movement disorder specialists^40^. Controls were self-reported free of neurological disease. Metadata were collected on over 40 variables including age, sex, race, geography, diet, medication, health, gastrointestinal issues, weight fluctuation, and body mass index. We enrolled 212 persons with PD and 136 controls in 2014 (dataset 1)^41^, and 323 PD and 184 controls during 2015–2017 (dataset 2)^11^. Subsequently, we excluded 11 PD and 4 control samples for failing 16S sequencing, 2 PD for unreliable metadata, and 15 controls for lacking genotypes from dataset 1; and 11 PD and 10 controls were excluded from dataset 2 for lacking genotype data. The sample size used in current analyses was 199 PD and 117 controls in dataset 1, and 312 PD and 174 controls in dataset 2 (Supplementary Table 1).

### Microbiome data

Methods for collection, processing and analysis of microbiome data have been reported in detail^11^, and raw sequences are publicly available at NCBI SRA BioProject ID PRJNA601994. Each subject provided a single stool sample at a single time point, and each sample was measured once. Briefly, for both datasets uniformly, DNA/RNA-free sterile cotton swabs were used to collect stool, DNA was extracted using MoBio extraction kits, and 16S rRNA gene hypervariable region 4 was sequenced using the same primers, but in two laboratories, resulting in 10x greater sequencing depth in dataset 2 than dataset 1. Sequences were demultiplexed using QIIME2 (core distribution 2018.6)^42^ for dataset 1 and BCL2FASTQ (Illumina, San Deigo, CA) for dataset 2. Bioinformatics processing of sequences was performed separately for each dataset, but using an identical pipeline (see Wallen et al^11^ for step-by-step protocol). Unique amplicon sequence variants (ASVs) were identified using DADA2 v 1.8^43^ and given taxonomic assignment using DADA2 and SILVA (v 132) reference database. Analyses were performed at genus/subgenus/clade level (here, referred to as taxa). Taxa that were associated with PD were then investigated at species level. This was important because not all species of *Corynebacterium_1*, *Porphyromonas*, and *Prevotella* are opportunistic pathogens. Species that made-up each taxon were identified by SILVA when an ASV matched a species at 100% homology. To augment SILVA, we blasted ASVs that made up *Corynebacterium_1*, *Porphyromonas*, and *Prevotella* against the NCBI 16S rRNA database for matches that were >99–100% identical with high statistical confidence.

### Genetic data

#### Defining SNCA region

Since expression of *SNCA* has been implicated in PD and the most significant genetic markers of PD map outside *SNCA* and are eQTL for *SNCA*, we set out to explore the entire region that includes known *cis-*eQTLs for *SNCA*. We used GTEx (V8 release) database and searched for eQTLs for *SNCA* (https://gtexportal.org/home/gene/SNCA). The search returned 1,749 entries which included 601 unique eQTLs. They span from ch4:90.6Mb at 5’ upstream *SNCA* to ch4:88.9Mb at 3’ downstream *SNCA* (GRCh38/hg38). We had genotypes for 2,627 SNPs in this region (excluding SNPs with MAF<0.1 and imputation quality score <0.8), and among them, we had captured 413 of the 601 eQTLs for *SNCA*. Interaction test was conducted for all 2,627 SNPs and the SNP with the highest interaction *P* value was chosen for genotype-stratified analysis.

Genotype data for the *SNCA* region were extracted from GWAS data. Since only some of the GWAS data have been published and most were generated recently and unpublished, we will provide the methods in detail. Dataset 1 is composed of a subset of the NGRC subjects who were genotyped in 2009 using Illumina HumanOmni1-Quad array (GWAS published in 2010)^36^ and were subsequently enrolled for microbiome study, and additional NGRC samples that were collected for microbiome studies in 2014 who were genotyped in 2018 using Illumina Infinium Multi-Ethnic array (unpublished data). Dataset 2 was enrolled into NGRC in 2015-2017 and genotyped in 2020 using Infinium Global Diversity Array (unpublished data). Genotyping and quality control (QC) of SNP genotypes are described below. Unless otherwise specified, QC was performed using PLINK 1.9 (v1.90b6.16)^44^.

#### HumanOmni1-Quad_v1-0_B BeadChip

Approximately 70% of subjects in dataset 1 (N=244) were genotyped in 2009 using the HumanOmni1-Quad_v1-0_B BeadChip for a GWAS of PD^36^, resulting in genotypes for 1,012,895 SNPs. Subjects were also genotyped using the Illumina Immunochip resulting in genotypes for 202,798 SNPs. QC of genotype data had been previously performed using PLINK v1.07,^36^ therefore, this process was redone for current study using an updated version of PLINK v1.9. The mean non-Y chromosome call rate for samples in both arrays was 99.9%. Calculation of identity-by-descent in PLINK using HumanOmni genotypes revealed no cryptic relatedness between samples (PI_HAT > 0. 15). A subset of SNP mappings were in NCBI36/hg18 build, and were converted to GRCh37/hg19 using the liftOver executable and hg18ToHg19.over.chain.gz chain file from UCSC genome browser (downloaded from https://hgdownload.soe.ucsc.edu/downloads.html). SNP filtering for both HumanOmni and Immunochip genotypes included removal of SNPs with call rate < 99%, Hardy-Weinberg equilibrium (HWE) *P* value < 1E-6, MAF < 0.01, and MAF difference between sexes > 0.15. HumanOmni and Immunochip data were then merged, and SNPs with significant differences in PD patient and control missing rates (*P* < 1E-5) and duplicate SNPs were removed. To remove duplicate SNPs, we first checked the genotype concordance between duplicated SNPs. If duplicate SNPs were concordant, we took the SNP with the lowest missing rate, or the first listed SNP if missing rates were the same. If duplicate SNPs were discordant, we removed both SNPs as we do not know which SNP is correct. After QC, the remaining number of genotyped SNPs was 910,083 with a mean call rate of 99.8%.

#### Infinium Multi-Ethnic EUR/EAS/SAS-8 Kit

Approximately 30% of subjects in dataset 1 (N=89) were enrolled after the 2010 PD GWAS. These samples were genotyped in 2018 using the Infinium Multi-Ethnic EUR/EAS/SAS-8 array. Raw genotyping intensity files were uploaded to GenomeStudio v 2.0.4 where genotype cluster definitions and calls were determined for each SNP using intensity data from all samples. The GenCall (genotype quality score) threshold for calling SNP genotypes was set at 0.15, and SNPs that resulted in a genotype cluster separation < 0.2 were zeroed out for their genotype. Genotypes for 1,649,668 SNPs were then exported from GenomeStudio using the PLINK plugin v 2.1.4, and converted to PLINK binary files for further QC. The mean non-Y chromosome call rate for samples was 99.8%. Calculation of identity-by-descent revealed no cryptic relatedness among samples (PI_HAT < 0.15). A subset of SNP mappings were in GRCh38/hg38 build, and were converted to GRCh37/hg19 using the liftOver executable and hg38ToHg19.over.chain.gz chain file. The same SNP filtering criteria were implemented here as described above for the first group in dataset 1: call rate < 99%, HWE *P* value < 1E-6, MAF < 0.01, MAF difference between sexes > 0.15, significant differences in PD patient and control missing rates (*P* < 1E-5), and removal of duplicate SNPs. After QC, the remaining number of genotyped SNPs was 749,362 with a mean call rate of 100%.

#### Infinium Global Diversity Array-8 v1.0 Kit

All subjects in dataset 2 (N=486) were genotyped at once in 2020 using the Infinium Global Diversity Array. Genotype clusters were defined using GenomeStudio v 2011.1 and 99% of the genotyped samples. Genotypes were not called for SNPs with GenCall score <0.15, and failure criteria for autosomal and X chromosome SNPs included the following: call rate < 85%, MAF ≤ 1% and call rate < 95%, heterozygote rate ≥ 80%, cluster separation < 0.2, any positive control replicate errors, absolute difference in call rate between genders > 10% (autosomal only), absolute difference in heterozygote rate between genders > 30% (autosomal only), and male heterozygote rate greater than 1% (X only). All Y chromosome, XY pseudo-autosomal region (PAR), and mitochondrial SNPs were manually reviewed. Genotypes for 1,827,062 SNPs were released in the form of PLINK binary files. The mean non-Y chromosome call rate for samples was 99.2%. Calculation of identity-by-descent showed two subjects were genetically related as a parent and offspring (PI_HAT = 0.5), which we were already aware of. The same SNP filtering criteria was implemented here as it was for dataset 1: call rate < 99%, HWE *P* value < 1E-6, MAF < 0.01, MAF difference between sexes > 0.15, significant differences in PD patient and control missing rates (*P* < 1E-5), and removal of duplicate SNPs. After QC, the remaining number of SNPs for dataset 2 was 783,263 with a mean call rate of 99.9%.

#### *Principal component analysis* (PCA)

We performed PCA for each genotyping array using 1000 Genomes Phase 3 reference genotypes. Study genotypes were first merged with 1000 Genomes Phase 3 genotypes (previously filtered for non-triallelic SNPs and SNPs with MAF > 5%) using GenotypeHarmonizer v 1.4.23^45^ and PLINK. Merged genotypes were then LD-pruned as previously described^36^, resulting in a mean LD-pruned subset of ~148,000 SNPs. Principal components were calculated using pruned SNPs and the top two PCs were plotted using ggplot2 (Supplementary Figure 1).

#### Imputation

To increase SNP density, we imputed genotypes using Minimac4^46^ on Trans-Omics for Precision Medicine (TOPMed) Imputation Server (https://imputation.biodatacatalyst.nhlbi.nih.gov)^47^. To be compatible with TOPMed, we converted SNP coordinates to GRCh38/hg38 using the liftOver executable and hg19ToHg38.over.chain.gz chain file. SNP mappings were then checked and corrected for use with TOPMed reference panels using the utility scripts HRC-1000G-check-bim.pl (v4.3.0) and CreateTOPMed.pl (downloaded from https://www.well.ox.ac.uk/~wrayner/tools/), and a TOPMed reference file ALL.TOPMed_freeze5_hg38_dbSNP.vcf.gz (downloaded from https://bravo.sph.umich.edu/freeze5/hg38/download). Running of these utility scripts resulted in a series of PLINK commands to correct genotypes files for concordance with TOPMed by excluding SNPs that did not have a match in TOPMed, mitochondrial SNPs, palindromic SNPs with frequency > 0.4, SNPs with non-matching alleles to TOPMed, indels, and duplicates. Once running of PLINK commands was complete, genotype files were converted to variant call format (VCF) and submitted to the TOPMed Imputation Server using the following parameters: reference panel TOPMed version r2 2020, array build GRCh38/hg38, *r*^2^ filter threshold 0.3 (although we excluded from downstream analyses SNPs with *r*^2^ <0.8), Eagle v2.4 for phasing, skip QC frequency check, and run in QC & imputation mode. VCF files with genotypes and imputed dosage data were then outputted by the imputation server and used in statistical analyses. Directly genotyped and imputed genotypes from HumanOmni1-Quad_v1-0_B BeadChip and Infinium Multi-Ethnic EUR/EAS/SAS-8 Kit arrays were merged to create dataset 1. To merge genotypes, one duplicate subject was first removed from the Infinium Multi-Ethnic array VCF files. Then, per chromosome VCF files were merged by first indexing the files using tabix, then merging the files using bcftools’ merge function (tabix and bcftools v 1.10.2). The genome-wide data included 20,263,129 SNPs (1,282,026 genotyped and 18,981,103 imputed) for dataset 1 and 21,389,007 SNPs (719,329 genotyped and 20,669,678 imputed) for dataset 2.

For the present study, the *SNCA* region was defined as ch4:88.9Mb-90.6Mb (as described above). SNPs within *SNCA* region with MAF<0.1 were excluded as there would be too few homozygotes for stratified analysis. Imputed SNPs with imputation quality score *r*^2^ <0.8 were also excluded. Analysis included 2,627 SNPs that were directly genotyped or imputed in both datasets.

#### Statistical analysis

For all analyses, raw taxa abundances were transformed using the centered log-ratio (clr) transformation before including in tests. The clr transformation was performed using the following formula in R:

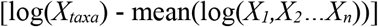

where *X_taxa_* is the raw abundance of either *Corynebacterium_1*, *Porphyromonas*, or *Prevotella* in a single sample with a pseudocount of 1 added, and *X_i_,X_2_…X_n_* are the raw abundances of every taxon detected in the same sample with a pseudocount of 1 added.

Throughout, tests were conducted in two datasets separately, and results were meta-analyzed using fixed- and random-effect models, and tested for heterogeneity. If heterogeneity was detected across two datasets (Cochran’s Q *P* < 0.1), random-effect meta-analysis results were reported. If no heterogeneity was detected (Cochran’s Q *P* ≥ 0.1), fixed-effect results were reported. *P* values were all two-tailed.

1. *Screening for Interaction*. We tested interaction to identify candidate SNPs that may modify the association of *Corynebacterium_1*, *Porphyromonas*, or *Prevotella* with PD. For each dataset separately, linear regression was performed using PLINK 2 (v2.3 alpha) --glm function to test the interaction between case/control status and SNP on the abundance of each taxon.

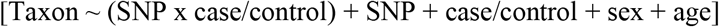

where taxon is the clr transformed abundance of *Corynebacterium_1*, *Porphyromonas*, or *Prevotella*, and SNP is genotype defined as dosages of the minor allele ranging from 0 to 2 in additive model. Interaction test was adjusted for sex, age, and main effects of case/control status and SNP. Interaction β and standard errors generated for each taxon were then used as input for meta-analysis in METASOFT v2.0.1^48^. Summary statistics are in Supplementary Tables 2-4. For each taxon, the SNP that reached the highest statistical significance in meta-analysis was tagged as candidate interacting SNP. *Linkage disequilibrium:* To visualize the results across the *SNCA* region, results from meta-analyses were uploaded to LocusZoom^49^. LD between SNPs was calculated in LocusZoom based on the “EUR” LD population. The resulting plots show the location of the SNPs tested in the region and their LD with candidate interacting SNP (Fig. 1a-c). To determine if the three candidate interacting SNPs were correlated, possibly tagging the same variant, or independent, pairwise LD estimates were calculated using the LDpair tool with 1000 Genome phase 3 European data from LDlink v4.1.^50^
2. *Association of taxa with PD as a function of genotype*. Subjects were grouped by their genotype at the interacting SNP. We used the best guessed genotype for the imputed SNPs and directly genotyped SNPs. Association of each taxon with PD (case/control status) was tested within each genotype, while adjusting for age and sex, using linear regression via the R function glm from the stats v 3.5.0 package. Odds ratios (OR) and corresponding *P* values were calculated using linear regression. Each dataset was analyzed separately. Meta-analysis was performed using the metagen function of the meta R package v4.9.7, specifying the summary measure to be “OR”. Results are shown in Table 1. Boxplots were created using ggplot2 v 3.1.0 (Fig. 2). Of the two variants of each SNP, the one that was associated with enhanced differential abundance in PD vs. controls was tagged as the effect allele.
3. *Association of interacting SNP with PD*. To test whether the interacting SNP had a main effect on PD risk, we used Firth’s penalized logistic regression (logistf R package v 1.23) testing SNP genotype (dosages of the effect allele ranging from 0 to 2) in an additive model against case-control status adjusting for age and sex. OR, SE and *P* values were calculated. Results were meta-analyzed using a fixed-effects model as implemented in the metagen function, of the meta R package v4.9.7, specifying the summary measure to be “OR”.

#### Functional analysis *in silico*

While we had defined the S*NCA* region such that it encompassed known eQTLs, only 413 of 2,676 SNPs tested were eQTL. Thus, if left to chance, the odds that a candidate SNP would be an eQTL was ~15%. We used UCSC Genome Browser (hg38 build) to map the candidate SNPs and visually inspect if they were in a regulatory sequence. To determine, for each SNP, if they were found in genome-wide studies to be significantly associated with gene expression, we used two eQTL databases, GTEx (https://gtexportal.org) and eQTLGen (https://www.eqtlgen.org).

## Data Availability

Individual-level raw 16S sequences and basic metadata are publicly available at NCBI Sequence Read Archive (SRA) BioProject ID PRJNA601994. Genetic data and summary statistics of interaction of 2,627 SNPs in *SNCA* region with PD on clr transformed abundances of taxa are provided in Supplementary Table 2 for *Coryenbacterium_1*, Supplementary Table 3 for *Porphyromonas*, and Supplementary Table 4 for *Prevotella*.

## Code availability

No custom codes were used. All software and packages, their versions, relevant specification and parameters are stated in the Methods section.

## Acknowledgements

This work was supported by the US Army Medical Research Materiel Command endorsed by the US Army through the Parkinson’s Research Program Investigator-Initiated Research Award under Award number W81XWH1810508 (to H.P.), National Institute of Neurological Disorders and Stroke grant R01 NS036960 (to H.P.), NIH Udall grants P50 NS062684 (to C.P.Z.) and P50 NS108675 (to D.G.S.), NIH Training Grant T32 NS095775 (to Z.D.W.) and NIH T32 GM008361 Medical Scientist Training Program (to W.J.S). Opinions, interpretations, conclusions, and recommendations are those of the authors and are not necessarily endorsed by the US Army or the NIH.

## Authorship contributions

Conception (HP), design (HP, ZDW, SAF, EM, CPZ, DGS), data acquisition (HP, SAF, EM, CPZ, DGS), data analysis (HP, ZDW, WJS), interpretation (HP, ZDW), drafting the manuscript (ZDW, HP) and revising it critically for important intellectual content (all authors).

## References

1 Collaborators, G. B. D. P. s. D. Global, regional, and national burden of Parkinson’s disease, 1990-2016: a systematic analysis for the Global Burden of Disease Study 2016. Lancet Neurol 17, 939–953, doi:10.1016/S1474-4422(18)30295-3 (2018).

2 Chang, D. et al. A meta-analysis of genome-wide association studies identifies 17 new Parkinson’s disease risk loci. Nat Genet, doi:10.1038/ng.3955 (2017).

3 Nalls, M. A. et al. Identification of novel risk loci, causal insights, and heritable risk for Parkinson’s disease: a meta-analysis of genome-wide association studies. Lancet Neurol 18, 1091–1102, doi:10.1016/S1474-4422(19)30320-5 (2019).

4 Tanner, C. M. Advances in environmental epidemiology. Mov Disord 25 Suppl 1, S58–62 (2010).

5 Hamza, T. H. et al. Genome-Wide Gene-Environment Study Identifies Glutamate Receptor Gene GRIN2A as a Parkinson’s Disease Modifier Gene via Interaction with Coffee. PLoS Genet 7, e1002237 (2011).

6 Cannon, J. R. & Greenamyre, J. T. Gene-environment interactions in Parkinson’s disease: specific evidence in humans and mammalian models. Neurobiol Dis 57, 38–46, doi:10.1016/j.nbd.2012.06.025 (2013).

7 Hill-Burns, E. M. et al. A genetic basis for the variable effect of smoking/nicotine on Parkinson’s disease. Pharmacogenomics J 13, 530–537, doi:10.1038/tpj.2012.38 (2013).

8 Biernacka, J. M. et al. Genome-wide gene-environment interaction analysis of pesticide exposure and risk of Parkinson’s disease. Parkinsonism Relat Disord 32, 25–30, doi:10.1016/j.parkreldis.2016.08.002 (2016).

9 Travagli, R. A., Browning, K. N. & Camilleri, M. Parkinson disease and the gut: new insights into pathogenesis and clinical relevance. Nat Rev Gastroenterol Hepatol 17, 673–685, doi:10.1038/s41575-020-0339-z (2020).

10 Horsager, J. et al. Brain-first versus body-first Parkinson’s disease: a multimodal imaging case-control study. Brain 143, 3077–3088, doi:10.1093/brain/awaa238 (2020).

11 Wallen, Z. D. et al. Characterizing dysbiosis of gut microbiome in PD: evidence for overabundance of opportunistic pathogens. NPJ Parkinsons Dis 6, 11, doi:10.1038/s41531-020-0112-6 (2020).

12 Schmidt, T. S. B., Raes, J. & Bork, P. The Human Gut Microbiome: From Association to Modulation. Cell 172, 1198–1215, doi:10.1016/j.cell.2018.02.044 (2018).

13 Morais, L. H., Schreiber, H. L. t. & Mazmanian, S. K. The gut microbiota-brain axis in behaviour and brain disorders. Nat Rev Microbiol, doi:10.1038/s41579-020-00460-0 (2020).

14 Fan, Y. & Pedersen, O. Gut microbiota in human metabolic health and disease. Nat Rev Microbiol 19, 55–71, doi:10.1038/s41579-020-0433-9 (2021).

15 Gerhardt, S. & Mohajeri, M. H. Changes of Colonic Bacterial Composition in Parkinson’s Disease and Other Neurodegenerative Diseases. Nutrients 10, doi:10.3390/nu10060708 (2018).

16 Boertien, J. M., Pereira, P. A. B., Aho, V. T. E. & Scheperjans, F. Increasing Comparability and Utility of Gut Microbiome Studies in Parkinson’s Disease: A Systematic Review. J Parkinsons Dis 9, S297–S312, doi:10.3233/JPD-191711 (2019).

17 Braak, H. et al. Staging of brain pathology related to sporadic Parkinson’s disease. Neurobiol Aging 24, 197–211 (2003).

18 Braak, H., Rub, U., Gai, W. P. & Del Tredici, K. Idiopathic Parkinson’s disease: possible routes by which vulnerable neuronal types may be subject to neuroinvasion by an unknown pathogen. J Neural Transm (Vienna) 110, 517–536, doi:10.1007/s00702-002-0808-2 (2003).

19 Shannon, K. M. et al. Alpha-synuclein in colonic submucosa in early untreated Parkinson’s disease. Mov Disord 27, 709–715, doi:10.1002/mds.23838 (2012).

20 Breen, D. P., Halliday, G. M. & Lang, A. E. Gut-brain axis and the spread of alpha-synuclein pathology: Vagal highway or dead end? Mov Disord 34, 307–316, doi: 10.1002/mds.27556 (2019).

21 Knudsen, K. et al. In-vivo staging of pathology in REM sleep behaviour disorder: a multimodality imaging case-control study. Lancet Neurol 17, 618–628, doi:10.1016/S1474-4422(18)30162-5 (2018).

22 Kim, S. et al. Transneuronal Propagation of Pathologic alpha-Synuclein from the Gut to the Brain Models Parkinson’s Disease. Neuron 103, 627–641 e627, doi:10.1016/j.neuron.2019.05.035 (2019).

23 Nalls, M. A. et al. Large-scale meta-analysis of genome-wide association data identifies six new risk loci for Parkinson’s disease. Nat Genet 46, 989–993, doi: 10.1038/ng.3043 (2014).

24 Mata, I. F. et al. SNCA variant associated with Parkinson disease and plasma alpha-synuclein level. Arch Neurol 67, 1350–1356 (2010).

25 Emelyanov, A. et al. SNCA variants and alpha-synuclein level in CD45+ blood cells in Parkinson’s disease. J Neurol Sci 395, 135–140, doi: 10.1016/j.jns.2018.10.002 (2018).

26 Consortium, G. Human genomics. The Genotype-Tissue Expression (GTEx) pilot analysis: multitissue gene regulation in humans. Science 348, 648–660, doi: 10.1126/science.1262110 (2015).

27 Tomlinson, J. J. et al. Holocranohistochemistry enables the visualization of alpha-synuclein expression in the murine olfactory system and discovery of its systemic anti-microbial effects. J Neural Transm (Vienna) 124, 721–738, doi:10.1007/s00702-017-1726-7 (2017).

28 Stolzenberg, E. et al. A Role for Neuronal Alpha-Synuclein in Gastrointestinal Immunity. J Innate Immun, doi:10.1159/000477990 (2017).

29 Mizuta, I. et al. YY1 binds to alpha-synuclein 3’-flanking region SNP and stimulates antisense non-coding RNA expression. J Hum Genet 58, 711–719, doi:10.1038/jhg.2013.90 (2013).

30 Elkouris, M. et al. Long Non-coding RNAs Associated With Neurodegeneration-Linked Genes Are Reduced in Parkinson’s Disease Patients. Front Cell Neurosci 13, 58, doi: 10.3389/fncel.2019.00058 (2019).

31 Villegas, V. E. & Zaphiropoulos, P. G. Neighboring gene regulation by antisense long non-coding RNAs. Int J Mol Sci 16, 3251–3266, doi:10.3390/ijms16023251 (2015).

32 Dunn, A. R. et al. Synaptic vesicle glycoprotein 2C (SV2C) modulates dopamine release and is disrupted in Parkinson disease. Proc Natl Acad Sci U S A, doi:10.1073/pnas.1616892114 (2017).

33 Chartier-Harlin, M. C. et al. Alpha-synuclein locus duplication as a cause of familial Parkinson’s disease. Lancet 364, 1167–1169, doi:10.1016/S0140-6736(04)17103-1 (2004).

34 Sulzer, D. et al. T cells from patients with Parkinson’s disease recognize alpha-synuclein peptides. Nature 546, 656–661, doi:10.1038/nature22815 (2017).

35 Schonhoff, A. M., Williams, G. P., Wallen, Z. D., Standaert, D. G. & Harms, A. S. Innate and adaptive immune responses in Parkinson’s disease. Prog Brain Res 252, 169–216, doi:10.1016/bs.pbr.2019.10.006 (2020).

36 Hamza, T. H. et al. Common genetic variation in the HLA region is associated with late-onset sporadic Parkinson’s disease. Nat Genet 42, 781–785 (2010).

37 Svensson, E. et al. Vagotomy and subsequent risk of Parkinson’s disease. Ann Neurol 78, 522–529, doi:10.1002/ana.24448 (2015).

38 Liu, B. et al. Vagotomy and Parkinson disease: A Swedish register-based matched-cohort study. Neurology 88, 1996–2002, doi:10.1212/WNL.0000000000003961 (2017).

39 Matheoud, D. et al. Intestinal infection triggers Parkinson’s disease-like symptoms in Pink1(-/-) mice. Nature 571, 565–569, doi:10.1038/s41586-019-1405-y (2019).

40 Gibb, W. R. & Lees, A. J. A comparison of clinical and pathological features of young-and old-onset Parkinson’s disease. Neurology 38, 1402–1406, doi:10.1212/wnl.38.9.1402 (1988).

41 Hill-Burns, E. M. et al. Parkinson’s disease and Parkinson’s disease medications have distinct signatures of the gut microbiome. Mov Disord 32, 739–749, doi: 10.1002/mds.26942 (2017).

42 Bolyen, E. et al. Reproducible, interactive, scalable and extensible microbiome data science using QIIME 2. Nat Biotechnol 37, 852–857, doi:10.1038/s41587-019-0209-9 (2019).

43 Callahan, B. J. et al. DADA2: High-resolution sample inference from Illumina amplicon data. Nat Methods 13, 581–583, doi:10.1038/nmeth.3869 (2016).

44 Chang, C. C. et al. Second-generation PLINK: rising to the challenge of larger and richer datasets. Gigascience 4, 7, doi:10.1186/s13742-015-0047-8 (2015).

45 Deelen, P. et al. Genotype harmonizer: automatic strand alignment and format conversion for genotype data integration. BMC Res Notes 7, 901, doi:10.1186/1756-0500-7-901 (2014).

46 Das, S. et al. Next-generation genotype imputation service and methods. Nat Genet 48, 1284–1287, doi:10.1038/ng.3656 (2016).

47 Taliun, D. et al. Sequencing of 53,831 diverse genomes from the NHLBI TOPMed Program. bioRxiv, 563866, doi:10.1101/563866 (2019).

48 Han, B. & Eskin, E. Random-effects model aimed at discovering associations in meta-analysis of genome-wide association studies. American journal of human genetics 88, 586–598, doi:10.1016/j.ajhg.2011.04.014 (2011).

49 Pruim, R. J. et al. LocusZoom: regional visualization of genome-wide association scan results. Bioinformatics 26, 2336–2337 (2010).

50 Machiela, M. J. & Chanock, S. J. LDlink: a web-based application for exploring population-specific haplotype structure and linking correlated alleles of possible functional variants. Bioinformatics 31, 3555–3557, doi:10.1093/bioinformatics/btv402 (2015).

