## Supplemental Figures for "Human-genome gut-microbiome interaction in Parkinson’s disease"

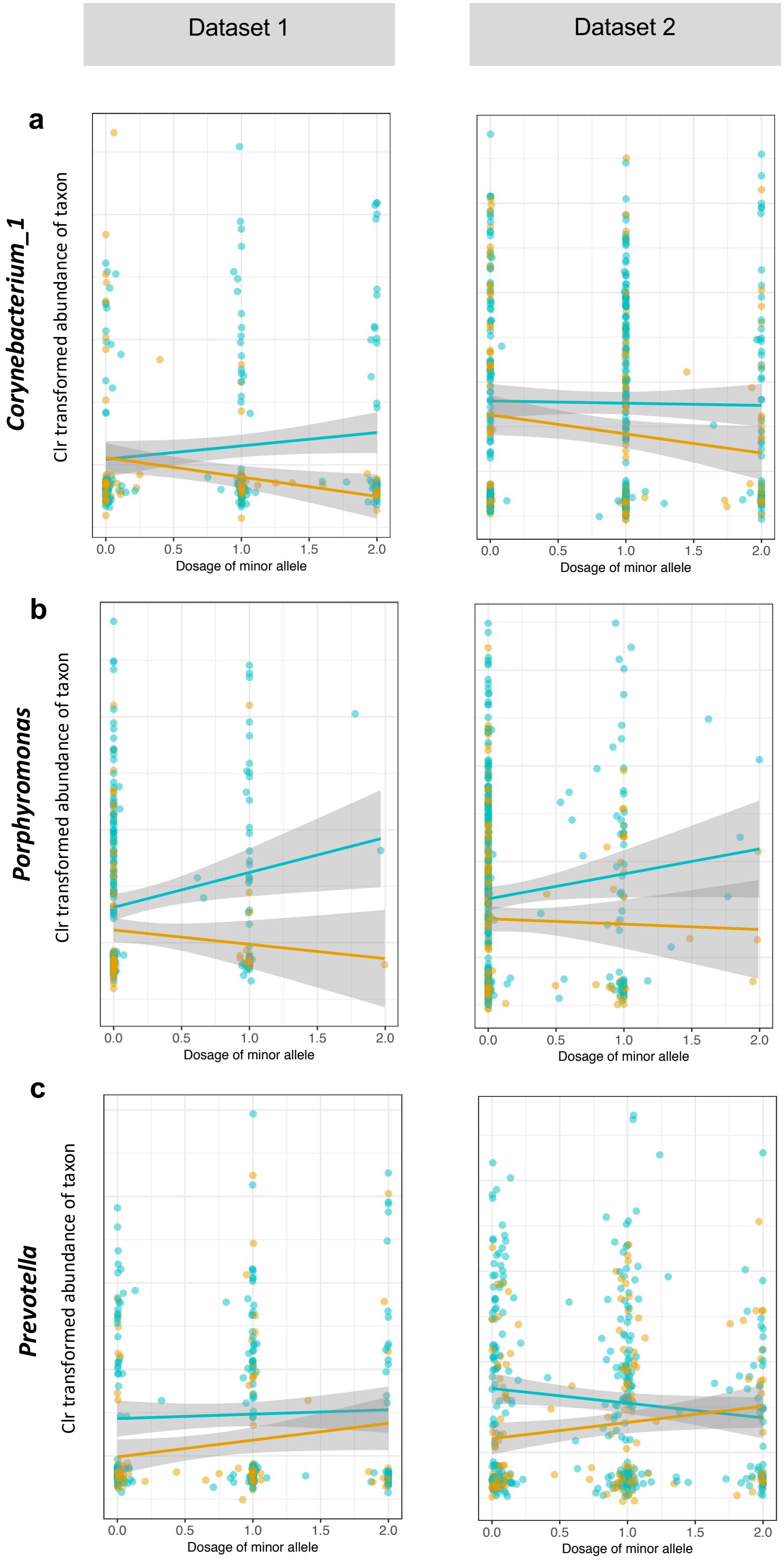


**Supplementary Fig. 1.** Visualizing the interaction of genotype, taxon, and case-control status through a plot of the raw data used in interaction scan. x axis: SNP genotype defined as estimated dosage of minor allele (ranging from 0 to 2 copies of the minor (less common) allele). y axis: clr-transformed abundance of the taxon. SNPs were imputed (imputation quality score >0.8). Orange: Controls. Blue: PD. (a) Effect of rs356229 genotype on the abundance of *Corynebacterium_1* in PD vs. control. (b) Effect of rs10029694 genotype on the abundance of *Porphyromonas* in PD vs control. (c) Effect rs6856813 genotype on the abundance of *Prevotella* in PD vs. control. Interaction was tested in each dataset separately, shown here side-by-side. N= 199 cases and 117 controls in dataset 1, 312 cases and 174 controls in dataset 2. Plots were generated using ggplot2 v3.1.0. Linear trendlines for case and control data points were overlayed onto plots using geom_smooth with the “lm” method from ggplot2.

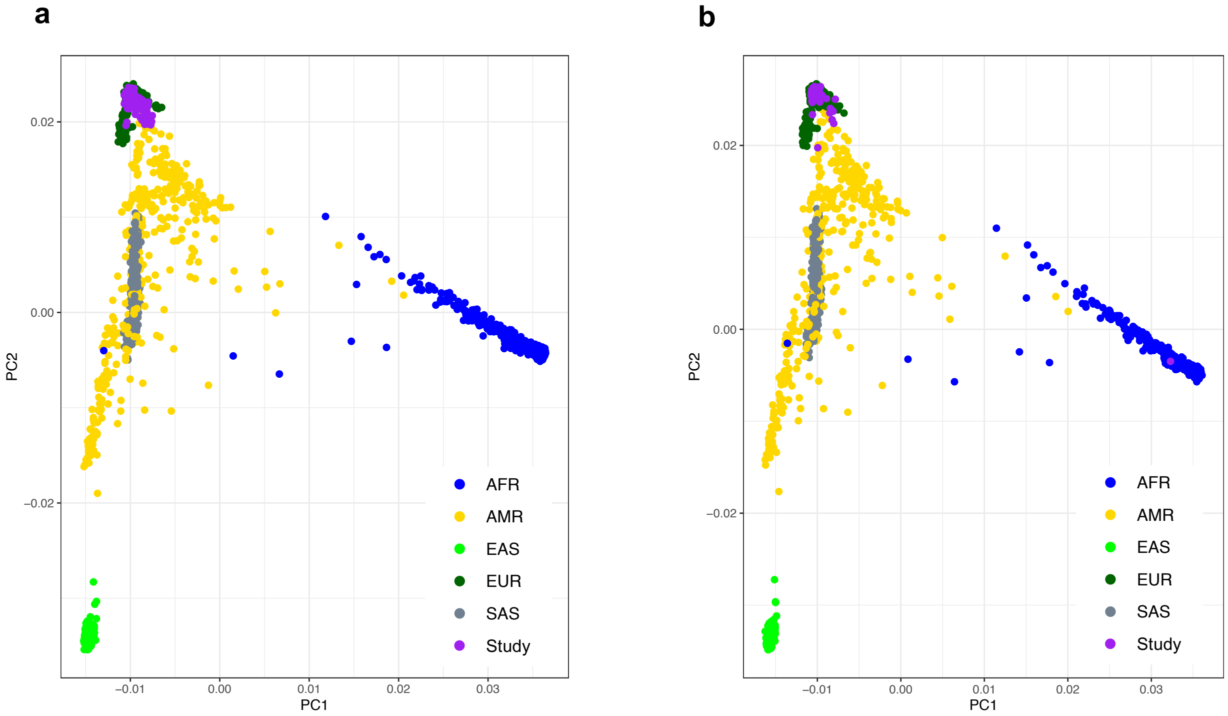


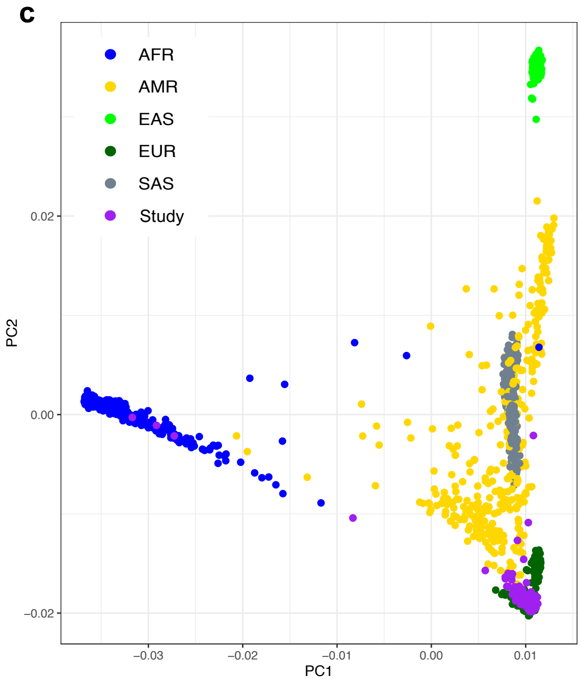


**Supplementary Fig. 2.**
Principal component analysis of study genotypes with 1000 Genome Phase 3 superpopulations. Principal component analysis was performed on merged, LD pruned study and 1000 Genome Phase 3 genotypes for samples genotyped using the HumanOmni1-Quad_v1-0_B BeadChip (a), Infinium Multi-Ethnic EUR/EAS/SAS-8 Kit (b), and Infinium Global Diversity Array-8 v1.0 Kit (c). All study subjects fall within a defined 1000 Genome Phase 3 superpopulation with no obvious outlying samples. AFR: African superpopulation from 1000 Genomes; AMR: Admixed American superpopulation from 1000 Genomes; EAS: East Asian superpopulation from 1000 Genomes; EUR: European superpopulation from 1000 Genomes; SAS: South Asian from 1000 Genomes; Study: study subjects genotypes.
